# Parallel processing relies on a distributed, low-dimensional cortico-cerebellar architecture

**DOI:** 10.1101/2022.07.10.499497

**Authors:** Eli J. Müller, Kevin Y. Hou, Fulvia Palesi, Joshua Tan, Thomas Close, Claudia A.M. Gandini Wheeler-Kingschott, Egidio D’Angelo, Fernando Calamante, James M. Shine

**Affiliations:** Complex Systems Research Group, The University of Sydney, Sydney, NSW, Australia; Brain and Mind Centre, The University of Sydney, Sydney, NSW, Australia; NMR Research Unit, Queen Square Multiple Sclerosis Centre, Faculty of Brain Sciences, UCL Queen Square Institute of Neurology, UCL, London, UK; Department of Brain and Behavioral Sciences, University of Pavia, Pavia, Italy; Brain Connectivity Research Center, IRCCS Mondino Foundation, Pavia, Italy; School of Biomedical Engineering, The University of Sydney, Sydney, NSW, Australia; Sydney Imaging, The University of Sydney, Sydney, NSW, Australia; National Imaging Facility, Sydney, NSW, Australia

## Abstract

A characteristic feature of human cognition is our ability to ‘multi-task’ – performing two or more tasks in parallel – particularly when one task is well-learned. How the brain supports this capacity remains poorly understood. Most past studies have focussed on identifying the areas of the brain – typically the dorsolateral prefrontal cortex – that are required to navigate information processing bottlenecks. In contrast, we take a systems neuroscience approach to test the hypothesis that the capacity to conduct effective parallel processing relies on a distributed architecture that interconnects the cerebral cortex with the cerebellum. The latter structure contains over half of the neurons in the adult human brain, and is well-suited to support the fast, effective, dynamic sequences required to perform tasks relatively automatically. By delegating stereotyped within-task computations to the cerebellum, the cerebral cortex can be freed up to focus on the more challenging aspects of performing the tasks in parallel. To test this hypothesis, we analysed task-based fMRI data from 50 participants who performed a task in which they either balanced an avatar on a screen (‘Balance’), performed serial-7 subtractions (‘Calculation’) or performed both in parallel (‘Dual-Task’). Using a set of approaches that include dimensionality reduction, structure-function coupling and time-varying functional connectivity, we provide robust evidence in support of our hypothesis. We conclude that distributed interactions between the cerebral cortex and cerebellum are crucially involved in parallel processing in the human brain.

## Introduction

How do distributed whole-brain neural activity patterns give rise to human cognitive function? This question lies at the heart of modern psychology and neuroscience but, despite decades of neuroimaging experiments, we still do not have a clear answer. One reason is that conventional neuroimaging methods applied to data from cognitive tasks typically represent the brain as a static snapshot of independent parts or at best, ‘functionally connected’ pairs of brain regions^1^. Another important issue is that neuroimaging experiments are usually designed to identify regions that are most selectively associated with a specific task, but are less well-suited to distinguishing the presence of multiple concurrent cognitive constructs within the same task^2^. For these reasons, many leading theories in cognitive neuroscience have relied on relatively static descriptions of the ‘key brain regions involved’ in a particular task.

In contrast to this view, there is evidence to suggest that the neural implementation of cognitive function in humans is far more dynamic and integrative^3^. In solving real world problems, we rarely isolate a specific cognitive capacity, such as focussed attention or resistance to distraction, but instead combine multiple cognitive constructs together in order to solve challenges in real-time^4^. Consider an experienced driver navigating heavy highway traffic in the pouring rain – the driver must remain focussed on the road, ensure the windshield wipers are on, regularly check their blind spots and also keep the pedals depressed at the appropriate level. This view of cognitive function in the real world is crucially dependent on the parallel processing of multiple distinct challenges, however for the reasons outlined above, we still lack a satisfying description of how the human brain is capable of supporting parallel processing.

To facilitate complex coordinated behavioural responses underpinned by similarly complex spatiotemporal activity patterns, the brain may first learn to execute at least one of the computations automatically (i.e., without paying close, conscious attention to the completion of the task). To achieve this, the system must be capable of responding to specific contexts with a high degree of spatial and temporal precision^5^. Secondly, the responses must be relatively error-free and reliable. Finally, the system must be able to be triggered in the presence of a specific stimulus or context without the need for deliberate attention. Without making the responses to different computational burdens relatively stereotyped in this fashion, performing two (or more) computations in parallel would require the prioritisation of one of the computations, likely to the detriment of the other task(s). In addition, any two tasks learned by the same network could potentially run into structural interference^6^, particularly if the networks required to complete the overlapping tasks use similar cortical regions.

Crucially, the architecture of the cerebellum is ideally suited to fulfil each of the features required for automatic processing, both in the sensorimotor and cognitive domains^7–10^. First, the cerebellum is organized in parallel modules with different cerebrocortical regions^8^. In direct contrast to the basal ganglia, the internal circuitry of the cerebellar cortex consists of sparse, distributed connectivity patterns that likely support dimensionality expansion^11^, rather than reduction (as is the case for the basal ganglia^12, 13^). In addition, the glutamatergic outputs of the cerebellum through the deep cerebellar nuclei innervate ‘core’ thalamic nuclei^14^, which project to the granular layers of the frontal cortex^15^ in a much more precise manner than the ‘matrix’ thalamic nuclei. There is also evidence that cerebellar circuits can condition on their own outputs, and hence learn to execute specific sequences of effects based on triggering context signals^16^. Anatomically, the cerebellum is bidirectionally interconnected with multiple cerebrocortical areas, with major tracts connecting the dentate nucleus to the frontal and prefrontal cerebral cortex, along with other associative areas^17, 18^. Functionally, the cerebellum plays a critical role in shaping complex functional network dynamics^19^, as evidenced by its role in both resting state^20, 21^ and task-related neuroimaging studies^7, 22–24^. Based on these architectural features and relationships with complex, dynamic neuroimaging patterns, we hypothesized that connections between the cerebellar cortex and cerebellum are crucial for the facilitation of parallel processing. Using a set of approaches that include dimensionality reduction, structure-function coupling and time-varying functional connectivity, we provide robust evidence in support of our hypothesis.

## Results

To test this hypothesis, we reanalysed an existing fMRI dataset^25^ consisting of 50 healthy individuals who performed a challenging ‘Dual-task’ in a 3T MRI scanner with their feet resting on a force-plate (Fig. 1A), and their vision oriented towards a 2-dimensional avatar that tilted forwards and backwards. There were three distinct trial types: during Balance blocks (Fig. 1B; blue), participants had to stabilize the slow fluctuations of the avatar using forward and backwards movements on the force-plate; during Calculation blocks (Fig. 1C; red), subjects had to track between 3-4 audible beeps, and then subtract that number, multiplied by 7, from a cue number presented prior to the trial; and during Dual-task blocks (Fig. 1D; purple), subjects performed both trials simultaneously.

**Fig 1.**
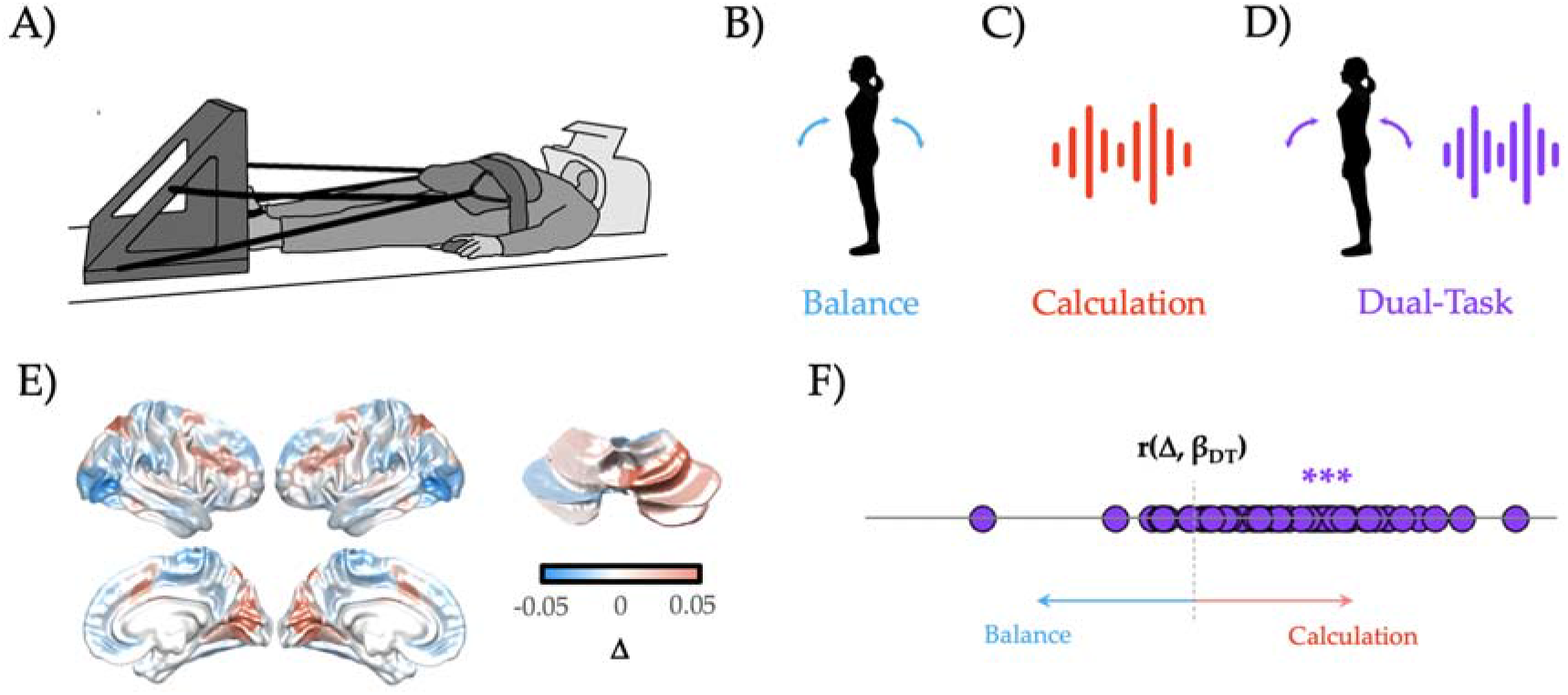
Low-dimensional balance between Integration and Segregation during Dual-task performance. A) participants lay supine in an MRI scanner, with their legs controlling a force-plate; B) Balance trials (blue) involved a dynamically moving avatar that the participant had to match; C) Calculation trials involved listening to a series of beeps, and then subtracting the multiple of 7 times the number of beeps from a cue number (red); D) Dual-task trials required performing both tasks, simultaneously (purple); E) The Calculation trials recruited increased BOLD in frontoparietal and visual cortices, along with right superior cerebellum, whereas Balance trials were associated with increased BOLD in lateral visual cortex, medial motor cortex and parietal operculum; F) the Dual-task β map across all 50 subjects was more similar to the Calculation β map (i.e., positive correlation with λ_1_) than the Balance β map (i.e., inverse correlation with λ_1_). *** - p < 0.001.

### Brain state signatures during dual-task performance

First, we compared the BOLD patterns associated with the performance of the three different tasks blocks. Specifically, we created a difference map between the average group-level β parameters estimated from 400 cortical and 28 cerebellar regions of interest for the Balance and Calculation trials (Δ). By comparing this difference map to the β map from the Dual-task trials – r(Δ,β_DT_) – we could determine whether performing the two tasks in tandem led to a brain map that was more or less like one or the other single tasks – a positive correlation with this map (λ_1_) was suggestive of the Dual-task reflecting the more challenging Calculation task; a negative correlation with the less challenging Balance task; and a null correlation with the notion of optimally splitting activity between the two (or a pattern orthogonal to the two single tasks). Consistent with the second option, we found that the low-dimensional signature of Dual-task performance was more similar to the Calculation β map than the Balance β map (r = 0.192 ± 0.05, p = 6.5x10^-5^; Fig. 1F), suggesting that during the Dual-task trials, the cerebral cortex and cerebellum configured their activity to ensure the effective completion of the Calculation trials.

Despite the brain states during Dual-task trials having more in common with the Calculation than the Balance trials, close examination of the RMS error of the Balance portion of the Dual-task trials suggests that subjects were performing the task as well as than when they performed the Balance trial on its own (Kolmogorov-Smirnov test: p = 0.358). So how was the brain configured on these Dual-task trials in order to mediate this stability? Based on previous empirical^23^ and theoretical^8, 26, 27^ work, we hypothesized that the distributed architecture integrating the cerebral cortex and cerebellum should be important for mediating this putative parallel processing performance. One straight-forward prediction is that balancing multiple tasks at the same time should recruit more regions of the cerebellum, and hence that cerebellar blood flow should be more strongly associated with Dual-Task performance than either the Balance or Calculation task alone. We found evidence to confirm this hypothesis – namely, a greater proportion of cerebellar regions were associated with a positive mean β value in Dual-Task as compared to Balance and/or Calculation trials (67.3% vs. 35.7% and 39.3%, respectively; χ^2^ (2, N = 50) = 249.6, p < 1.0x10^-4^).

### Unique patterns of cortico-cerebellar functional connectivity during dual-task performance

Given that the Dual-Task trials were more similar to Calculation trials than Balance trials (Figure 1F), how was the brain capable of supporting multiple tasks at the same time? We hypothesized that Balance, Calculation and Dual-task trials should have unique patterns of cortico-cerebellar functional connectivity that could allow the brain to support multiple channels of communication within the same system. To test this hypothesis, we calculated the time-varying functional connectivity between all cortical and cerebellar parcels using the Multiplication of Temporal Derivatives approach (window = 20 TRs^28^) and then contrasted the three trial types with one another. We observed robust differences between the three trial types (Fig. 2). For instance, Calculation trials (when compared to Balance trials) were associated with wide-spread cortico-cerebellar connectivity between lobule V and the majority of cortical networks, as well as more targeted connections between VIIIa/IX and primary sensorimotor networks (Fig. 2A). In contrast, Balance trials (when compared to Calculation trials) showed predominant increases in intermediate cerebellar lobules (e.g., Crus I and II) with higher-order cortical networks. In contrast, Dual-task trials were associated with heightened frontoparietal connections with intermediate cerebellar lobules, particularly Crus I and VIIIa, when compared to both Balance (Fig. 2B) and Calculation trials (Fig. 2C).

**Figure 2.**
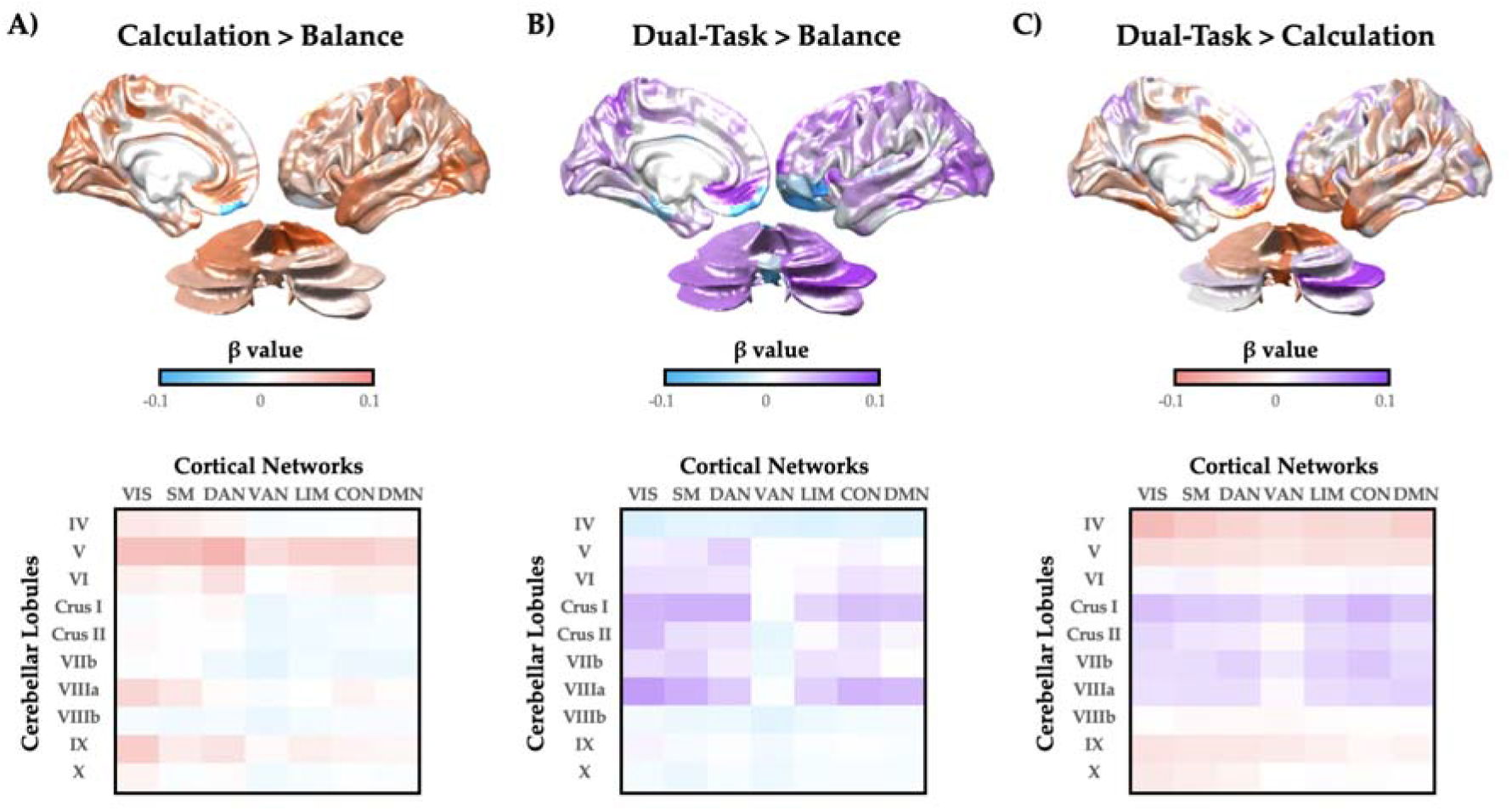
Cortico-cerebellar task-based functional connectivity. A) patterns of task-based cortico-cerebellar functional connectivity during Calculation (red) when compared to Balance (blue) trials – upper: mean task-based connectivity strength for cerebral cortex and cerebellum; lower: mean task-based connectivity strength collapsed into 7 Yeo networks (columns) and 10 average lobules (rows); B) similar maps for Dual-task (purple) versus Balance; C) similar maps for Dual-task versus Calculation. Note: results were consistent for left and right hemispheres. Key: VIS – visual; SM – somatomotor; DAN – dorsal attention network; VAN – ventral attention network; LIM – limbic network; CON – control network; DMN – default mode network.

Having confirmed a robust relationship between the cerebral cortex and cerebellum during Dual-task performance, we next asked whether cortico-cerebellar functional connectivity patterns differentiated between correct and error Dual-task trials. To test this hypothesis, we fit a General Linear Model to each Dual-task trial, independently, for each cortico-cerebellar time-varying connectivity score. We then split Dual-task trials into correct (accurate calculation and small RMS error [< 50% of population distribution]) and incorrect (inaccurate calculation, large RMS error [>50%] or both) trials and compared (using a set of independent-samples T-tests) the task-based functional connectivity between cortical and cerebellar parcels as a function of effective Dual-task performance. We conducted a permutation test (5,000 iterations) to determine the likelihood of each edge being distinct between the two groups by chance. To summarize these results, we computed the mean significant β-value for the functional connectivity between each cerebellar lobule (averaged across hemispheres, and ignoring the connections of the vermis; from the cerebellar SUIT atlas^29^) and each of 7 pre-identified cortical networks (the Yeo 7 parcellation from the 400-region Schaefer atlas^30^; Fig 3). We found a robust increase in task-based functional connectivity between the ventral attention network (VAN) and lobules Crus II, VIIb, VIIIa and VI (Fig. 3), as well as more distributed connections between lobule X and multiple cortical sub-networks. In contrast, Crus I was relatively functionally disconnected from all cortical networks (except VAN) during effective dual-task performance, which is consistent with known patterns of cerebellar lesion-related cognitive impairments^31^.

**Fig. 3.**
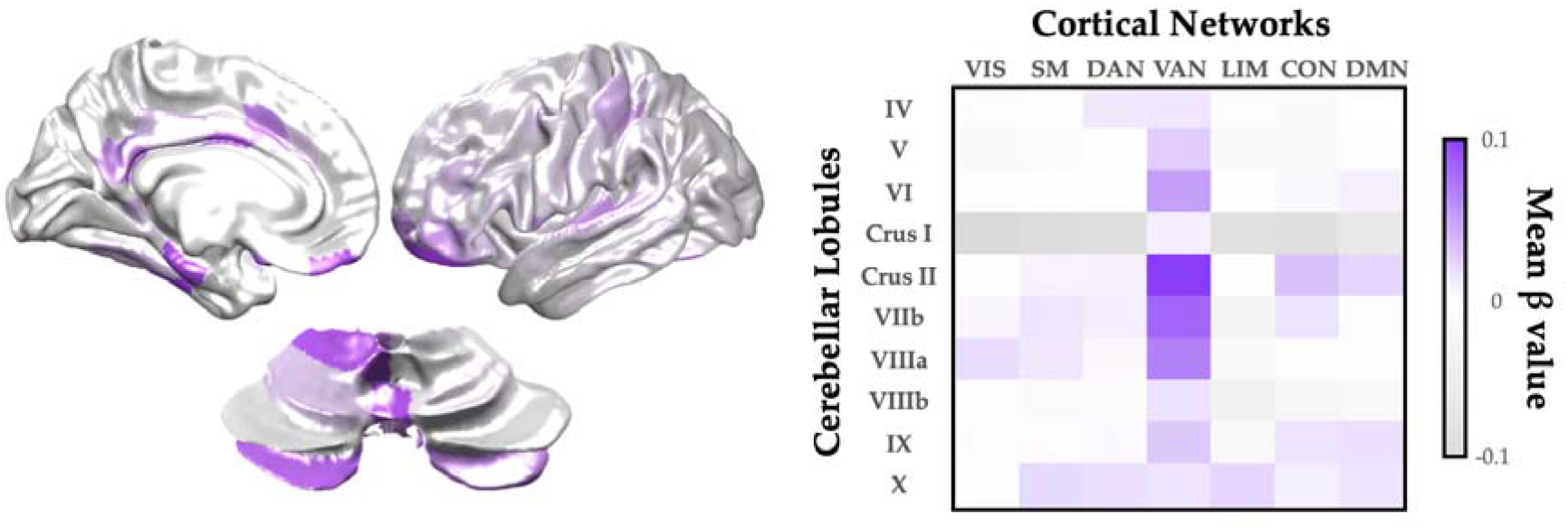
Increased cortico-cerebellar task-based functional connectivity associated with successful Dual-Task performance. Left: Key cortical and cerebellar regions with heightened task-based functional connectivity during Dual-task trials with correct vs. incorrect answers. Right: Mean significant β-value (cortical sub-network [Yeo 7 atlas] vs. cerebellar lobule [SUIT atlas]) associated with task-based functional connectivity values for correct vs. incorrect Dual-Task performance (p < 0.001; random permutation test). Key: VIS – visual; SM – somatomotor; DAN – dorsal attention network; VAN – ventral attention network; LIM – limbic network; CON – control network; DMN – default mode network.

### Dual-task performance balances network integration and segregation

One way in which the distributed cortico-cerebellar architecture could facilitate effective parallel processing is by striking an effective balance between integration and segregation^32–34^. In previous work, we have used a combination of time-varying functional connectivity and a topological measure that quantifies network-level integration – the Participation Coefficient (PC)^35^ – to demonstrate that the systems-level network structure of functional connectivity changes during task performance, with cognitively challenging tasks requiring higher integration than relatively simple tasks^35^. From this, we predicted that the Balance task should be relatively segregated (i.e., low PC), the Calculation task should be relatively integrated (i.e., high PC) and the Dual-task trials should strike a balance between the two extremes (i.e., intermediate PC). Using our standard time-varying analysis (see Methods), we observed robust evidence for our predictions (Fig. 4; F_2,147_ = 3.41; p = 0.036). In addition, although the Dual-task topological pattern was positively correlated with the average of Balance and Calculation (r = 0.464; p < 0.001), it was not a direct super-position of the two maps, suggesting topological reconfiguration during the different task states. Together, these results confirm that parallel processing in the brain is supported by a topological balance between integration and segregation.

**Figure 4.**
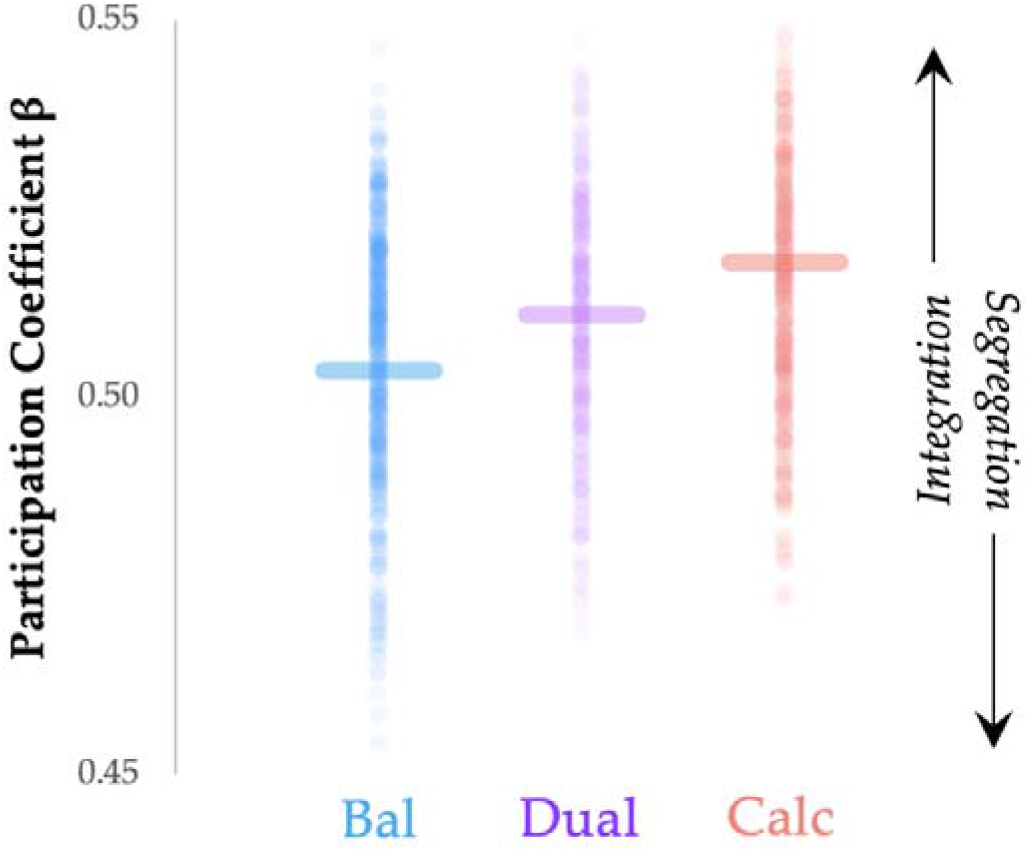
Parallel processing balances integration and segregation. Balance trials were associated with relative Segregation (low PC; blue), Calculation trials with relative Integration (high PC; red) and Dual-task trials with a balance between Integration and Segregation (intermediate PC; purple) – F_2,147_ = 3.41; p = 0.036. Thick lines represent the median value for each group.

### Cortico-cerebellar activity flow mapping

The input and output streams of cerebral cortex and cerebellum interact via distinct white matter pathways. Importantly, while the structural connections between these two structures are reciprocal, they are imbalanced^17, 18^ – different pathways exist from the cortex to the cerebellum (and *vice versa*). Specifically, thick-tufted layer V pyramidal neurons in the deep layers of the cerebral cortex send projections to the mossy fibre pathway of the cerebellum (via the pontine nuclei), thus forming the Cortico-Ponto-Cerebellar (CPC) tract (Fig. 5A). In contrast, the cerebral cortex receives feedback from the cerebellum via the deep cerebellar nuclei, which project via the ‘Core’ nuclei of the thalamus – i.e., the Cerebello-Thalamo-Cortical (CTC) tract (Fig. 5B). Plastic changes between the mossy fibre pathway and the Purkinje cells of the cerebellar cortex are proposed to act as a major site for the refinement of automatized behaviour^10, 26, 27, 36^ and hence, the capacity to perform multiple tasks simultaneously. From our observations that the timeseries of the cerebral cortex and cerebellum were highly coordinated during Dual-task behaviour, we hypothesized that the specific patterns of BOLD activity in both the cortex and cerebellum should be related to the intersection between prior BOLD activity in the cerebellum (via the CTC) and cerebral cortex (via the CPC).

**Fig. 5.**
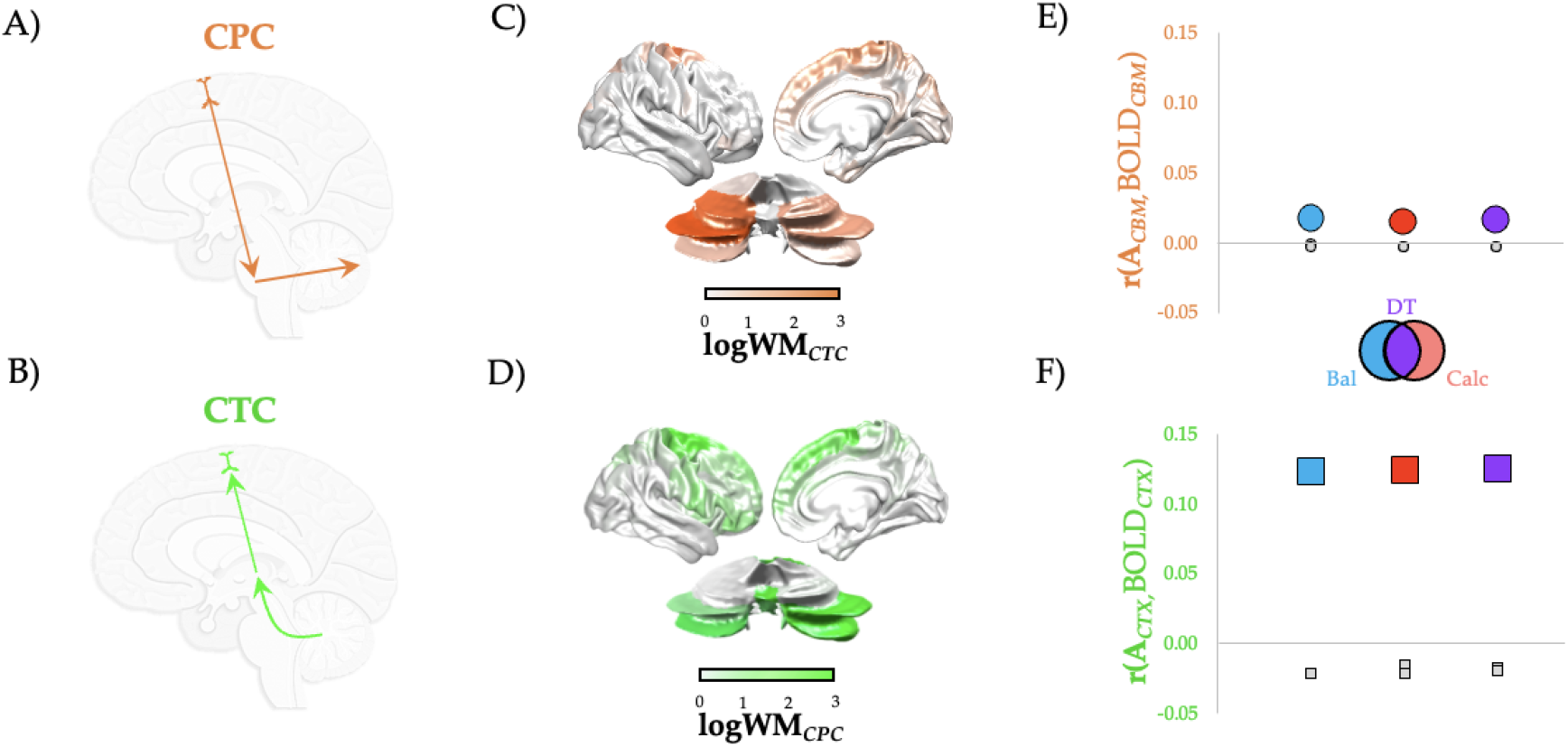
Cortico-cerebellar Structure-Function Mapping across Trial types. Two different major tracts connecting the cerebral cortex and cerebellum: the Cortico-Ponto-Cerebellar (CPC; orange; A) tract sends projections from the cortex via the pontine nuclei into the mossy fibres of the cerebellum, and the Cerebello-Thalamo-Cortical (CTC; green; B) tract derives from the deep cerebellar nuclei, which project back via the core thalamic nuclei to the cerebral cortex; C) normalized (in log_10_ of white matter connectivity) map of projections from the cerebral cortex to cerebellum via CPC (orange); D) normalized (in log_10_ of white matter connectivity) map of projections from cerebellum to the cerebellar cortex via CTC (green); E) activity flow mapping^37^ between cerebellar BOLD patterns predicted from CPC tract in Balance (Bal; blue), Calculation (Calc; orange) and Dual-Task (DT; purple) trials (circles) – see Methods for details; F) the same for cortical BOLD patterns predicted from the CTC tract (squares). All activity flow map correlations were greater than permuted null levels.

To test this hypothesis, we adapted the activity flow mapping approach^37^ to incorporate the structural connectivity between the cortex and cerebellum. Specifically, we extracted structural connectivity weights for both the contralateral CPC (Fig. 5C; orange) and CTC (Fig. 5D; green) tracts^17^ from a cohort of 28 healthy individuals from the Human Connectome Project^17^ (i.e., not the same individuals that performed the parallel processing task). While both tracts are over-expressed in the frontal cortices, there were relatively more CPC projections from the parietal lobes and more CTC projections that innervate the frontal cortex, which is consistent with known anatomical projection patterns^8, 26, 38, 39^. A parsimonious interpretation of these data is that the frontal cortex benefits from the information provided to the cerebellum by posterior cortices that process potential opportunities for action (also known as affordances^40^).

If cortico-cerebellar communication is required for effective Dual-task performance, then blood flow within either cortex or cerebellum during Dual-task trials should be predictive of subsequent blood flow (assuming sufficient delay) within the cortical (or cerebellar) regions to which they are connected by white matter projections. To create an estimate of what these predicted BOLD responses should be, we created two template maps – one for predicted cerebellar activity (estimated cerebellar activity: *A_CTX_* = *W_CBM_*. *CPC*) and one for predicted cortical activity (estimated cortical activity: *A_CBM_* = *W_CTX_*. *CTC*) – by multiplying the cortico-cerebellar structural connectivity matrices with the pre-processed BOLD pattern observed during the three different trial types. We then correlated these prediction vectors with the actual BOLD patterns in the respective regions. If the observed patterns of activity were similar, we can conclude that BOLD activity patterns were intimately related to the reciprocal structural connections between the cerebral cortex and cerebellum.

Across all three trial types, both cortico-cerebellar (via CPC; Fig. 5E, circles) and cerebello-cortical (via CTC; Fig. 5F, squares) activity flow patterns were significantly greater for matched vs. un-matched data (all p < 0.05), suggesting that functional activity was coordinated by connections both from the cerebral cortex to the cerebellum (i.e., CPC) and vice versa (i.e., CTC) across all tasks. Interestingly, despite the consistent positive relationships, cerebello-cortical connections (i.e., CTC) were more robustly able to predict subsequent cortical patterns than cortico-cerebellar connections (i.e., CPC), suggesting that the feedback from the cerebellum to the cerebral cortex was more crucial for task performance. Finally, we found that the match between A_CTX_/A_CBM_ and the raw data was greater in correct versus incorrect Dual-task trials for both cerebral cortex (T = 2.397, *p* = 0.017) and cerebellum (T = 2.049, *p* = 0.041), further confirming the importance of cortico-cerebellar interaction for parallel processing.

## Discussion

In this study, we used systems-level neuroimaging analysis to demonstrate that robust interactions between the cerebral cortex and cerebellum are associated with effective Dual-task performance. We hypothesized that, through distributed white matter pathways that interconnect these major cortical systems, the brain can differentiate different task contexts so as to effectively maintain the performance of two computational tasks in parallel. To test this hypothesis, we analysed BOLD data from the cerebral cortex and cerebellum, and in doing so demonstrated that Dual-Task performance recruited heightened cerebellar activity (Figure 1) and functional connectivity between the cerebral cortex and cerebellum (Figures 2 and 3) that was linked to the balance between integration and segregation (Figure 4) and related to the structural connections between the cerebellum and cerebral cortex (Figure 5). Together, these results highlight the importance of systems-level interactions in the manifestation of complex cognitive capacities.

Our results clearly demonstrate that models that incorporate the cerebellum and its massive, high-dimensional architecture provide a more parsimonious account for how the brain can balance the challenges inherent with parallel processing^8, 26^. The connectivity between the cortex and cerebellum is optimally set-up to fulfil this capacity. Specifically, the major output of the cerebral cortex – layer V PT-type pyramidal neurons – provides the primary afferent input to the cerebellar cortex (i.e., granule cells), by way of the pontine nuclei^8, 26, 41^. Following a massive dimensionality expansion that has been argued to facilitate pattern separation^11^, the outputs of the cerebellum (the deep cerebellar nuclei) send large glutamatergic projections to the ventral tier of the thalamus^39^, wherein they innervate the cerebral cortex. The thalamic targets of the cerebellum then go on to drive activity in the cerebral cortex, typically in a high-frequency, precise fashion^42^ that we have argued to form the basis of relatively automatic modes of behaviour^26, 27^. Here, we extend these functional neuroanatomical principles to incorporate the completion of challenging, dual tasks. We anticipate that similar patterns will be observed in future experiments that interrogate different types of dual tasks, particularly those in which one (or both) of the tasks is capable of relative automaticity. Whether such automaticity benefits extend to purely perceptual tasks, such as the attentional blink^43^, is an interesting open question for future work.

The topological signature of functional networks estimated from BOLD data have previously been linked to effective performance on cognitive tasks. For instance, an integrated brain has been linked to the completion of a range of complex tasks, such as those that probe working memory^35, 44, 45^, logical reasoning^46^ and attentional tracking^47, 48^. In contrast, a relatively segregated functional network has been linked to relative sensorimotor automaticity^32, 33^, as well as to attentional vigilance^49^. Our results are consistent with the spectrum implied by these previous results – the Balance task, which presumably tapped into relatively well-learned behaviours, was associated with a segregated functional network; and the Calculation task, which likely required more focussed, flexible attention, was associated with a relatively integrated network. Interestingly, although the Dual-task trials were arguably more challenging than the Calculation trials on their own, the topology of the network actually demonstrated a balance between integration and segregation, suggesting that performing tasks in parallel requires an ability to avoid topological extremes, perhaps so as to maximise information processing capabilities^50^. In addition, there are theoretical reasons to believe that the finite nature of biological networks may imbue specific limits on the number of possible tasks that can be run in parallel, although we expect that the high-dimensional architecture of the cerebellum^11^ will likely boost this capacity, particularly as a function of experience^26, 27^. Precisely which systems in the brain help to control this balance remains an open question, however there is intriguing results to suggest that the neuromodulatory system may play a crucial role in this process^26, 51, 52^.

Systems-level neuroimaging analysis provides an integrated perspective of cognitive capacities^33^, however BOLD dynamics are necessarily indirect – i.e., they don’t measure neural activity directly, but rather filtered through the low-dimensional lens of perfusion^53, 54^. While BOLD signal remains a robust measurement for neural signalling^55, 56^, it only reveals a part of how the brain functions. This is particularly true for the cerebellar cortex, whose complex, convoluted anatomy^8, 57^ and idiosyncratic firing properties^16, 58, 59^ render simple, linear readouts of neural activity from BOLD problematic. Specifically, there is evidence to suggest that BOLD measurements in the cerebellar cortex predominantly track activity in the mossy fibre pathway (via the CPC)^60, 61^, whereas outputs from the Purkinje cells (via the CTC) are more difficult to characterize with BOLD signalling^62, 63^. While this does suggest caution with respect to the interpretation of our results, it makes the presence of robust cerebello-cortical activity flow mapping via the CTC (Figure 5F) all the more fascinating of a result, as it suggests that the fate of the Purkinje cells is relatively sealed by the specific pattern of mossy fibre inputs that they received, although we anticipate that this mapping is likely augmented by the process of learning – i.e., it should be less profound when facing highly novel task contexts. Irrespective, we hope that by consolidating analysis from multiple neuroimaging techniques that we have provided a robust illustration of changes to cortico-cerebellar circuits during a parallel processing task.

The capacity to perform tasks in parallel clearly scales positively with experience. In the future, it will be fascinating to examine the interactions between the cerebral cortex and cerebellum as individuals learn to perform individual tasks to relative automaticity. There is robust empirical previous work linking cerebellar output with highly-overtrained behaviours in rodents^64^. Similar arguments have been made when analysing automaticity in the performance of challenging cognitive tasks^23^. Interestingly, there is also evidence suggesting that, over the course of learning a simple sensorimotor task, the brain shifts from a relatively integrated to a segregated architecture^32, 33^. This suggests a novel prediction: the extent to which a particular task has been well-learned will lead to relative segregation of the topological network signature of the brain, which in turn will make the task easier to automatise, and hence to combine successfully with other, more challenging dual tasks.

## Conclusions

Here, we have demonstrated that dynamic interactions between the cerebral cortex and cerebellum are critically related to the performance of a challenging dual-task. Future research is required to determine whether similar principals are related to parallel processing of other simultaneous cognitive and perceptual challenges, as well as across distinct spatiotemporal scales.

## Methods

### Experimental setup

The functional data from this study arose from a re-analysis of a previously published dataset^25^ – here, we will include the minimal information required to interpret the results, and point the interested reader to the original study for full details. Participants lay supine in the MRI scanner with their feet against a custom-made force platform attached to the MRI bed (Fig 1A; sample frequency of 100 Hz), with the position of the force platform was adjusted to subject height. A four-button device was placed underneath the right hand for the calculation task. The tasks were projected onto a white screen placed at the head of the scanner. Participants could see the screen via a mirror attached to the head coil.

During the Balance task, an avatar in the shape of a woman was displayed on the screen. The avatar swayed forward and backward. Participants were instructed to try to keep the avatar in the upright position by increasing or decreasing the level of plantar flexion force measured by the load cell. As in normal standing, increasing the plantar flexion force led to a backward sway (and v.v.). At the start of every balance condition, participants were given two seconds to bring the avatar in the upright position. After these two seconds, a disturbance signal was added, causing the avatar to sway forward and backward. To keep the avatar upright, participants had to counteract these disturbances. The disturbance signal was made by combining fifteen sinusoidal signals with random phases and with frequency characteristics based on an average frequency spectrum of centre of pressure movement during upright standing (0.025–1 Hz), measured in ten young and ten old adults. The maximum amplitude of the disturbance was ±30°. The error for each Balance trial was created by calculating the sum of the Root Mean Squared error between the optimal balanced avatar (i.e., 90^0^) and the position of the actual avatar. Trials were subsequently median split to identify ‘good’ and ‘bad’ balance trials.

The Calculation task consisted of serial subtractions with increments of seven – at the beginning of each trial, a number between 50 and 100 was projected on the screen for two seconds, after which a plus sign was displayed on the screen and a beep was generated every 3 to 4 seconds through an MRI compatible headphone (MR confon Optime 1, Magdeburg, Germany), with a total of four beeps per trial. Participants were instructed to subtract the number ‘7’ with every beep. At the end of each trial, four answer possibilities were displayed on the screen: one indicating the correct answer, two erroneous answers, and the option that none of the other answers is correct. Participants indicated which answer they thought was correct by pressing the corresponding button of the four-button device.

During the Dual-task condition, subjects performed the Balance and Calculation tasks simultaneously. The distribution of RMS errors in the Balance trials and Dual-task trials were compared using a Kolmogorov-Smirnov test.

An fMRI block-design was used to alternate between the three conditions: Balance, Calculation, and Dual-task. Every participant performed twelve blocks, each block including one trial of each condition (thus three trials), with the order of the conditions randomized, both across blocks and between subjects. At the end of every block a 15-second rest period was given in which the participants fixated their gaze on a plus sign.

### MRI acquisition and pre-processing

Brain imaging was performed on a 3-T SIEMENS Magnetom Skyra System (Siemens, Erlangen, Germany) with a 20-channel head/neck coil. For functional scans, a T2*-weighted multiband gradient echo-planar imaging (EPI) sequence was used (TR = 700 msec, TE = 30 msec, flip angle = 55°, 48 axial slices, slice thickness = 3 mm, no gap, in-plane resolution 3x3 mm)^65^. After the functional scanning session, a high-resolution magnetization prepared rapid acquisition gradient echo (MPRAGE) T1-weighted sequence (TR = 2,100 msec, TE = 4.6 msec, TI = 900 msec, flip angle = 8°, 192 contiguous slices, voxel resolution 1 mm³, FOV = 256x256x192 mm, iPAT factor of 2) was obtained in sagittal orientation. These anatomical scans were used to co-register the functional runs using SPM 12. The anatomical scan was segmented using the SPM tissue probability maps. All functional scans were co-registered to the anatomical scan and normalized to the Montreal Neurological Institute (MNI) template brain via the forward deformations revealed by the segmentation.

### Brain parcellation

Following pre-processing, the mean time series was extracted from 400 pre-defined cortical parcels using the Schaefer atlas^30^ and 28 pre-defined cerebellar parcels from the SUIT atlas^29^ (cerebellar nuclei were not included). The mean BOLD signal intensity from each region was extracted and then used for subsequent analyses.

### General Linear Model and Principal Component Analysis

A general linear model was fit to pre-processed, parcellated BOLD data with separate terms modelling each trial type (i.e., Balance, Calculation and Dual-task). The proportion of cerebellar regions associated with positive cerebellar β-values was compared across Balance, Calculation and Dual-task trials using a χ^2^ test with degrees of freedom = (rows – 1) x (columns – 1) = (3-1) x (2-1) = 2.

The average β-value for the Balance and Calculation trials were demeaned and analysed with a Principal Component Analysis. The coefficient of the leading principal component was correlated with the mean β map from the Balance and Calculation trials to demonstrate its utility as a linear decoder between Balance and Calculation. The dot-product between the Dual-task β map for each subject and the leading principal component was calculated, and then subjected to a 1-sample t-test to determine whether the loading was more similar to Calculation (positive loadings) or Balance (negative loadings).

### Time-varying functional connectivity

We used the multiplication of temporal derivatives (MTD) approach^28^ to calculate time-resolved dynamic functional connectivity between the selected ROIs; code is freely available at https://github.com/macshine/coupling/) with a window size of 20 TRs. For each node, n, with time points, t, a vector of t-1 temporal derivatives was calculated and normalized (temporal derivatives divided by the standard deviation of temporal derivatives, σ). Then, we created a matrix of functional coupling between the *i^th^* and *j^th^* nodes for each time point, by multiplying the temporal derivatives of each pair of nodes across each time point.

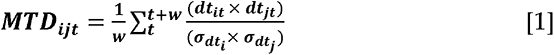

where *dt* is the first temporal derivative of the *i^th^* and *j^th^* time-series, and σ standard deviation of the temporal derivative, w is the window length of the simple moving average^28^. The MTD values for the cortico-cerebellar system (i.e., 400 x 28 = 11,200 edges) were entered into a similar General Linear Model to the cortico-cerebellar BOLD values, with a permutation test (5,000 iterations) used to test for statistical significance.

### Modularity Maximization

The Louvain modularity algorithm from the Brain Connectivity Toolbox (BCT^66^; http://www.brain-connectivity-toolbox.net) was used on the neural network edge weights to estimate community structure. The Louvain algorithm iteratively maximizes the modularity statistic, *Q*, for different community assignments until the maximum possible score of *Q* has been obtained (see Equation 2). The modularity of a given network is therefore a quantification of the extent to which the network may be subdivided into communities with stronger within-module than between-module connections.

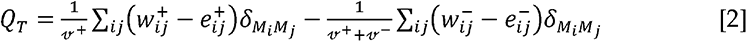

where *ν* is the total weight of the network (sum of all negative and positive connections), *w_ij_* is the weighted and signed connection between regions *i* and *j*, *e_ij_* is the strength of a connection divided by the total weight of the network, and *δ_MiMj_* is set to 1 when regions are in the same community and 0 otherwise. ‘+’ and ‘–‘ super-scripts denote all positive and negative connections, respectively.

For each epoch, we assessed the community assignment for each region 500 times and a consensus partition was identified using a fine-tuning algorithm (BCT). We calculated all graph theoretical measures on un-thresholded, weighted and signed connectivity matrices^66^. The stability of the γ parameter was estimated by iteratively calculating the modularity across a range of γ values (0.5-2.5; mean Pearson’s r = 0.859 +-0.01) on the time-averaged connectivity matrix for each subject – across iterations and subjects, a γ value of 1.0 was found to be the least variable, and hence was used for the resultant topological analyses.

### Participation Coefficient

The participation coefficient, PC, quantifies the extent to which a region connects across all modules (i.e., between-module strength) and has previously been used to successfully characterize hubs within brain networks^7, 35^. The PC for each region was calculated within each temporal window using Equation 3, where k_isT_ is the strength of the positive connections of region i to regions in module s at time *T*, and k_iT_ is the sum of strengths of all positive connections of region *i* at time *T*. Negative connections were discarded prior to calculation. The participation coefficient of a region is therefore close to 1 if its connections are uniformly distributed among all the modules and 0 if all of its links are within its own module.

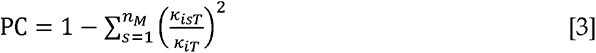

The participation coefficient for each parcel was compared across Balance, Calculation and Dual-task trials using paired t-tests (FDR *q* = 0.05).

### Diffusion MRI Analysis

The minimally processed HCP diffusion datasets (which included correction for motion, susceptibility distortions, gradient non-linearity and eddy currents) were subject to additional image processing, which multi-shell multi-tissue constrained spherical deconvolution to generate the fibre orientation distribution (FOD) in each voxel^67–69^. These steps were implemented in accordance with previous work^70^ and were performed using the MRtrix software package (http://www.mrtrix.org^71, 72^).

The T1-weighted images were used to generate a so-called ‘five-tissue-type’ (5TT) image^73^ using FSL^74^; the 5TT image classifies the voxel into one of 5 tissue types: cortical grey matter, sub-cortical grey matter, white matter, cerebrospinal fluid, and ‘5^th^ type’ (e.g., pathology). The FOD data and the 5TT image were used to generate 120 million streamlines using the anatomically constrained tractography (ACT) framework^73^, using dynamic and the 2^nd^-order Integration over Fibre Orientation Distributions (iFOD2)^71^ probabilistic fibre-tracking algorithm, using default MRtrix parameters, with the exception of FOD cutoff 0.06, maximum length 250 mm, step size 1 mm, and backtrack specified. This set of streamlines is referred to as the *whole-brain-tractogram* thereafter.

The cerebello-thalamo-cortical (CTC) and cortico-ponto-cerebellar (CPC) tracts were extracted from the *whole-brain-tractogram* by using contralateral cerebral and cerebellar cortices, cerebellar peduncles, contralateral red nuclei, and thalami as regions of interest (see ^17, 18^ for more details). To define the strength of the cerebellar connectivity with each of brain parcel, the log_10_ of the number of streamlines was used to weight the CTC and CPC tracts^75, 76^.

### Cortico-cerebellar activity flow mapping

To determine whether cortico-cerebellar interactions could transform cortical or cerebellar task-evoked activity into respective cerebello-cortical task activity, we modified the activity flow mapping procedure^37^ to incorporate estimates of cortico-cerebellar (CPC) and cerebello-cortical (CTC) structural connectivity. Specifically, for each trial type, block and subject, we calculated:

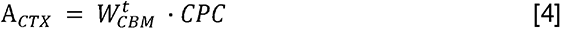

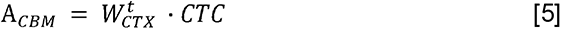

where *W_t_* is the evoked response estimate for every cortical (W*_CTX_*) or cerebellar (W*_CBM_*) parcel, CPC and CTC are the structural connectivity matrices described above, and *A_CTX_* and *A_CBM_* are the predicted activity pattern for each subgroup. For each trial type, block and subject, the predicted cortical and cerebellar activity patterns were then empirically compared to the observed activity patterns using Pearson correlations. A series of t-tests were used to compare the Pearson’s correlation loadings, with the non-matching predictions (e.g., using the cortical BOLD for Balance trials to predict cerebellar BOLD for Calculation trials) used a simple null model that contained all the same spectral features but spatiotemporal sequences that did not match the data. Finally, we created separate null distributions following a random permutation^77^ of both CPC and CTC, separately (each with 5,000 iterations).

## Acknowledgements

Data were provided by the Human Connectome Project, WU-Minn Consortium (PI: David Van Essen and Kamil Ugurbil; 1U54MH091657) funded by the 16 NIH Institutes and Centers that support the NIH Blueprint for Neuroscience Research; and by the McDonnell Center for Systems Neuroscience, Washington University.

JMS was supported by the National Health and Medical Research Council (GNT1193857). FC was supported by the National Health and Medical Research Council (GNT1117724) and the Australian Research Council (DP170101815). ED, CGW-K and FP received funding from H2020 Research and Innovation Action Grants Human Brain Project (#945539, SGA3). ED also received funding from the MNL Project “Local Neuronal Microcircuits” of the Centro Fermi (Rome, Italy). CGW-K receives funding from the MS Society (#77), Wings for Life (#169111), Horizon2020 (Human Brain Project), UCL-UCLH Biomedical Research Center (London, UK) BRC (#BRC704/CAP/CGW), MRC (#MR/S026088/1), Ataxia UK. CGWK is a shareholder in Queen Square Analytics Ltd. CGW-K is also a shareholder in Queen Square Analytics. TC was supported by the National Imaging Facility, a National Collaborative Research Infrastructure Strategy (NCRIS) capability.

---

**Figure S1.**
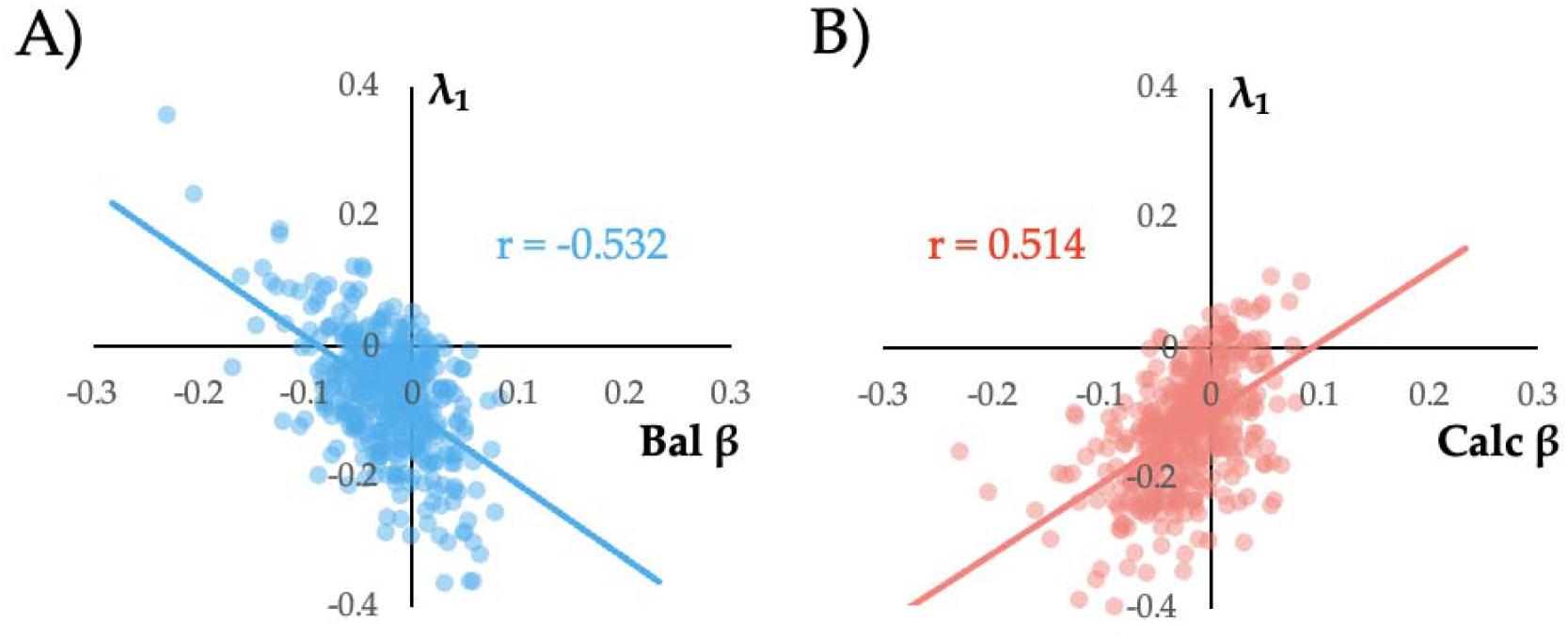
Correlation between Balance and Calculation maps and leading principal component. A) Loadings of the principal eigenvector (λ_1_) in both cerebral cortex (left) and cerebellum (right); F) scatter between mean Balance β map and λ_1_ (r = -0.532; p*_SPIN_* = 0.002); G) scatter plot between the mean Calculation β map and λ_1_ (r = 0.514; p*_SPIN_* < 0.0001).

